## Supplementary Information for "Transcranial focused ultrasound stimulation of the anterior temporal lobe enhances semantic memory by modulating brain morphology, neurochemistry and neural dynamics"

Behavioural results

Supplementary Figure 1

Supplementary Figure 2

Supplementary Table 1

Supplementary Table 2

Supplementary Table 3

Supplementary Table 4

Behavioural results

A 2 x 2 repeated measures ANOVA was conducted with stimulation (ATL vs. ventricle) and session (PRE vs. POST) as a within-subject factors to evaluate the effects of tbTUS in each task.

In the semantic task, there was a significant main effect of session (F _1,21_ = 10.61, p = 0.004) and an interaction between the stimulation and session (F _1,21_ = 5.09, p = 0.035) on accuracy. *Post hoc* t-tests revealed that ATL tbTUS significantly increased accuracy in semantic task (t= -3.66, p < 0.001). No significant effects were found in the control task (Fs < 1.73, ps > 0.202).

For reaction time (RT), we only found a significant main effect of session in both tasks (semantic: F _1,21_ = 88.56, p < 0.001; control: F _1,21_ = 8.56, p = 0.008). *Post hoc* t-tests showed that participants responded faster in the post-session, regardless of stimulation type or task (ATL stimulation – semantic: t = 11.13, p < 0.001; Ventricle stimulation – semantic: t = 6.39, p < 0.001; ATL stimulation – control: t = 2.37, p = 0.027; ventricle stimulation – control: t = 3.05, p = 0.006).

**Supplementary Figure 1**

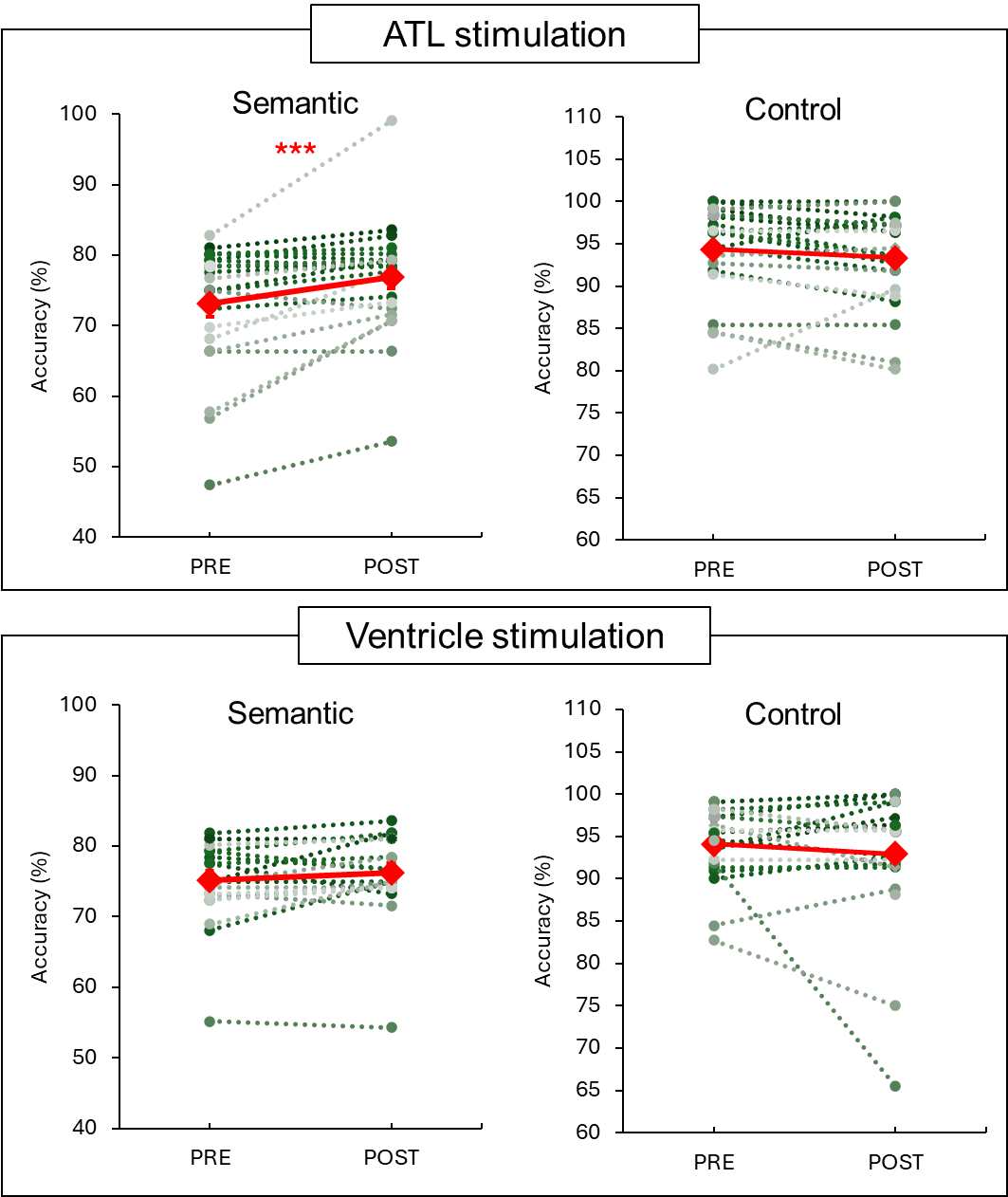

**Figure S1.** Task performance: accuracy. Red diamonds represent the mean of data. Circles represent each individual data. *** p < 0.001

**Supplementary Figure 2**

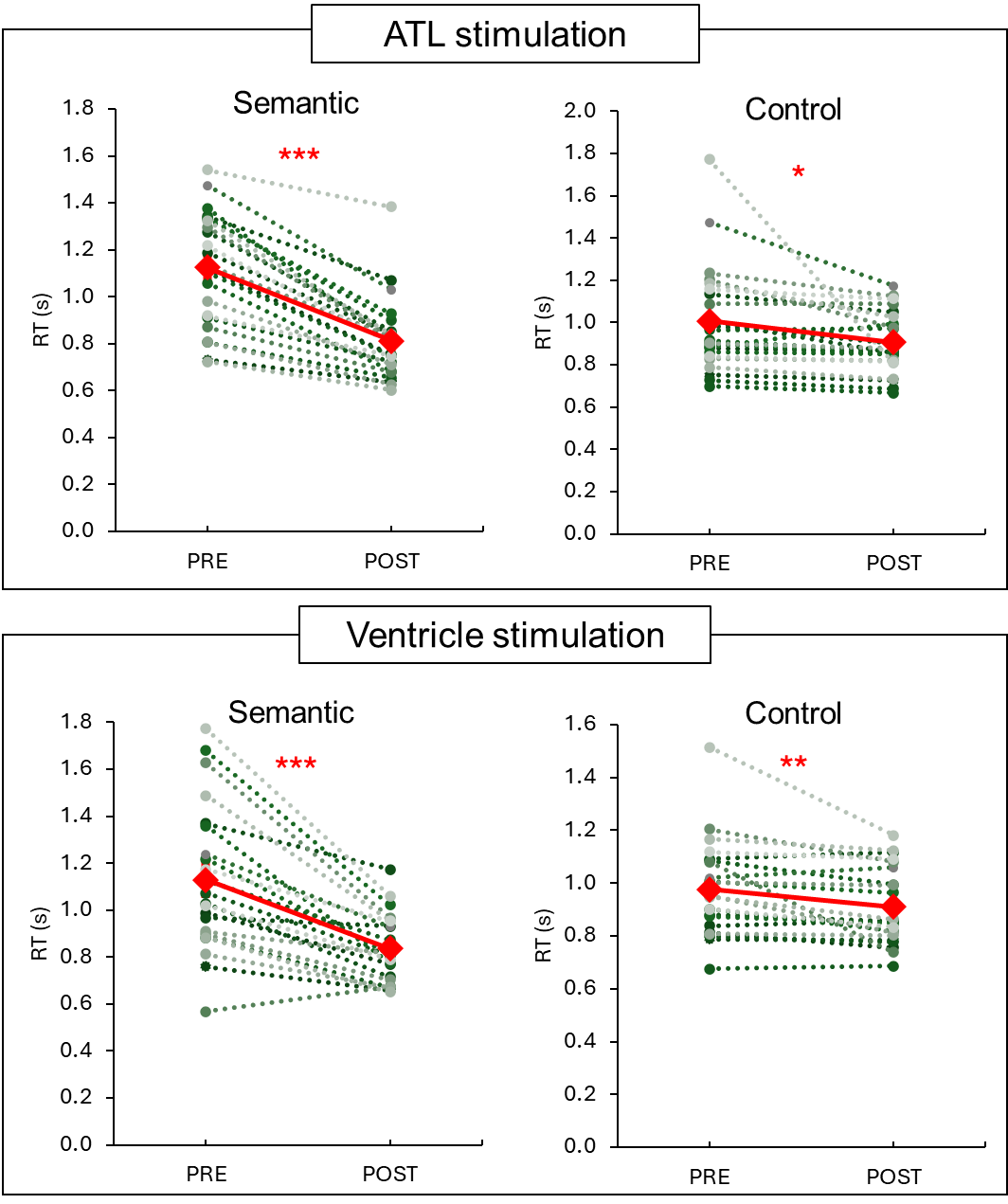

**Figure S2.** Task performance: RT. Red diamonds represent the mean of data. Circles represent each individual data. *** p < 0.001, ** p < 0.01, * p < 0.05

**Supplementary Figure 3**

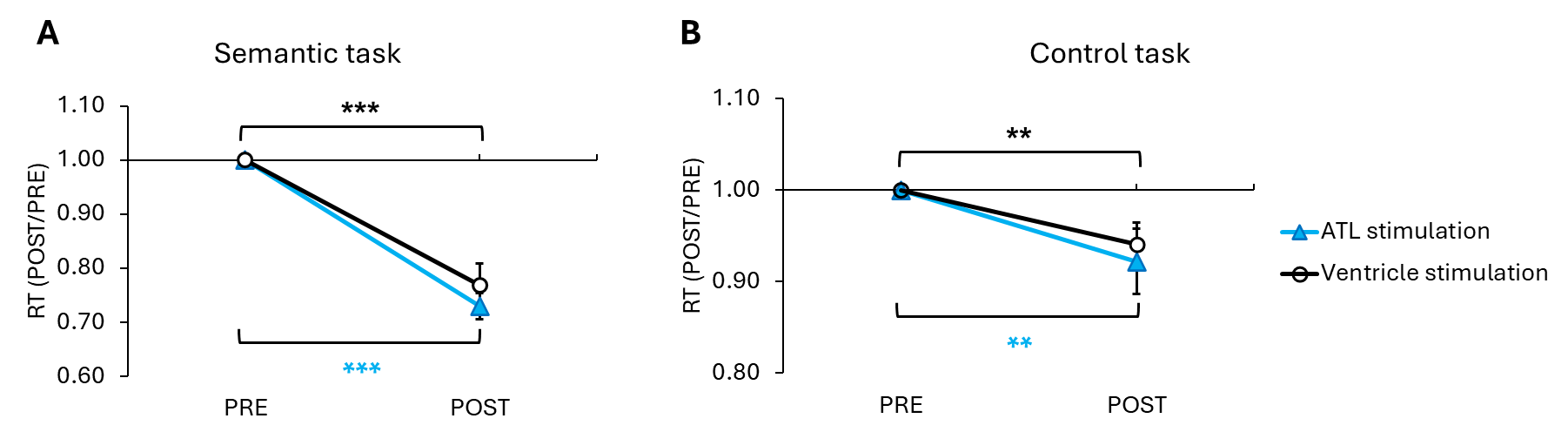

Semantic task

RT (POST/PRE)

|  | ATL stimulation | Ventricle stimulation |
| --- | --- | --- |
| PRE | 1 | 1 |
| POST | 0.729839 | 0.768347 |

**Figure S3.** A) tbTUS-induced changes in the normalised RT in the semantic task. B) tbTUS-induced changes in the normalised RT in the control (pattern matching) task. Light blue lines indicate the ATL stimulation. Black lines represent the control (ventricle) stimulation. Error bars represent standard error. *** p < 0.001, ** p < 0.01

**Supplementary Figure 4**

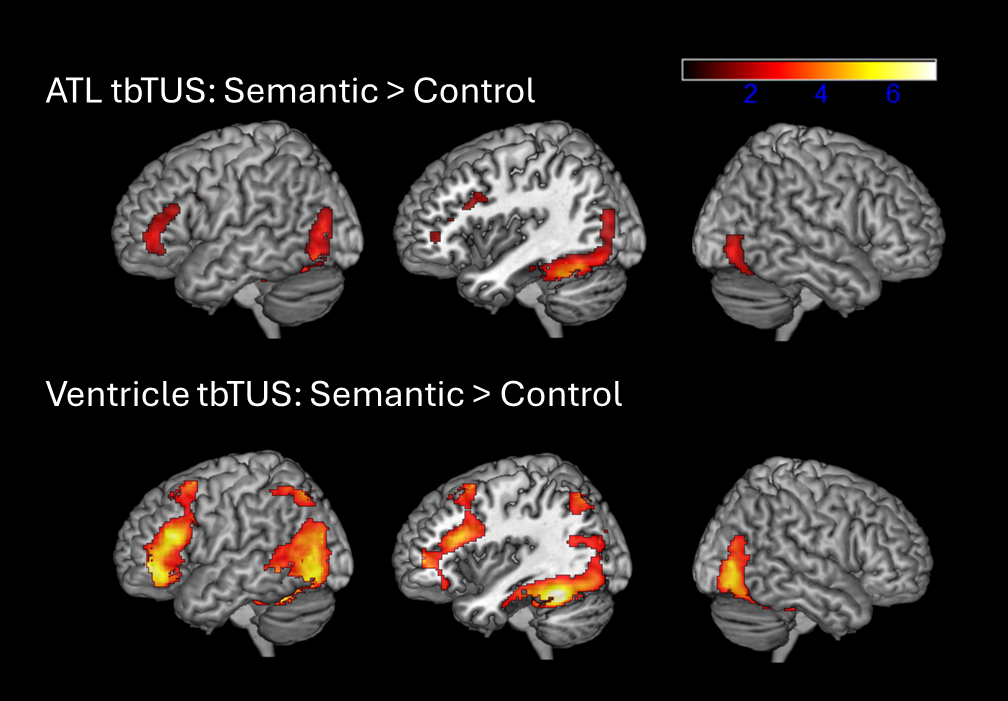

**Figure S4.** The result of fMRI for the contrast of Semantic > Control. The colour bar indicates T score.

**Supplementary Table 1**

|  |  |  | Data quality metrics | | | | | | | | | |  | Tissue segmentations | | |
| --- | --- | --- | --- | --- | --- | --- | --- | --- | --- | --- | --- | --- | --- | --- | --- | --- |
| Region | sonication | | | N | Linewidth (FWHM) | SNR | Fit error (%) | | | | | |  | GM (%) | WM (%) | CSF (%) |
|  |  | | |  | Water | Water | Water | GABA+ | glx | NAA | tCr | cho |  |  |  |  |
| ATL | ATL tbTUS | | | 20 | 23.96 (4.62) | 25314 (8255) | 0.74 (0.35) | 6.67 (1.19) | 5.30 (1.60) | 1.40 (0.50) | 2.31 (0.72) | 4.68 (0.72) |  | 0.49 (0.04) | 0.49 (0.04) | 0.02 (0.01) |
|  | Ventricle tbTUS | | | 19 | 28.57 (7.79) | 20763 (6143) | 0.78 (0.42) | 8.24 (3.33) | 6.01 (2.66) | 1.35 (0.78) | 2.12 (0.89) | 4.79 (0.87) |  | 0.49 (0.03) | 0.48 (0.03) | 0.02 (0.02) |
| OCC | ATL tbTUS | | | 17 | 10.97 (1.03) | 86252 (24501) | 0.45 (0.10) | 2.58 (0.56) | 2.31 (0.44) | 1.01 (0.34) | 1.68 (0.23) | 6.73 (0.72) |  | 0.69 (0.03) | 0.23 (0.03) | 0.07 (0.03) |
|  | Ventricle tbTUS | | | 21 | 10.93 (1.39) | 80140 (16955) | 0.43 (0.89) | 2.47 (0.54) | 2.09 (0.50) | 1.93 (0.37) | 1.60 (0.19) | 6.76 (0.71) |  | 0.68 (0.03) | 0.24 (0.03) | 0.07 (0.03) |

Table S1. The summary of MRS data quality metrics and tissue segmentation information. Data are shown as the mean (SD).

**Supplementary Table 2**

|  | ATL stimulation | | | |  | Ventricle (control) stimulation | | | |  | ATL stimulation vs. Ventricle stimulation | |
| --- | --- | --- | --- | --- | --- | --- | --- | --- | --- | --- | --- | --- |
| Parameter |  |  |  |  |  |  |  |  |  |  |  | |
| Intrinsic connectivity | Mean | SD | t | p |  | Mean | SD | t | p |  | t | p |
| L.ATL → R.ATL | 0.088 | 0.028 | 15.284 | <0.001 |  | 0.015 | 0.003 | 26.663 | <0.001 |  | 12.490 | <0.001 |
| L.ATL → L.IFG | 0.104 | 0.027 | 18.742 | <0.001 |  | 0.014 | 0.002 | 32.692 | <0.001 |  | 16.513 | <0.001 |
| L.ATL → R.IFG | 0.102 | 0.039 | 12.545 | <0.001 |  | 0.014 | 0.002 | 34.313 | <0.001 |  | 10.843 | <0.001 |
| L.ATL → L.pMTG | 0.062 | 0.040 | 7.535 | <0.001 |  | 0.016 | 0.002 | 30.652 | <0.001 |  | 5.790 | 0.013 |
| L.ATL → R.pMTG | 0.041 | 0.044 | 4.409 | <0.001 |  | 0.016 | 0.002 | 33.959 | <0.001 |  | 2.696 | <0.001 |
| R.ATL → L.ATL | 0.122 | 0.040 | 14.642 | <0.001 |  | 0.015 | 0.003 | 26.771 | <0.001 |  | 12.570 | <0.001 |
| R.ATL → L.IFG | 0.081 | 0.033 | 11.878 | <0.001 |  | 0.015 | 0.002 | 39.399 | <0.001 |  | 10.145 | <0.001 |
| R.ATL → R.IFG | 0.091 | 0.040 | 10.975 | <0.001 |  | 0.014 | 0.002 | 41.191 | <0.001 |  | 9.433 | <0.001 |
| R.ATL → L.pMTG | 0.093 | 0.043 | 10.513 | <0.001 |  | 0.016 | 0.002 | 37.783 | <0.001 |  | 8.760 | <0.001 |
| R.ATL → R.pMTG | 0.086 | 0.055 | 7.592 | <0.001 |  | 0.016 | 0.002 | 42.359 | <0.001 |  | 6.101 | <0.001 |
| L.IFG → L.ATL | 0.132 | 0.034 | 18.674 | <0.001 |  | 0.015 | 0.002 | 32.672 | <0.001 |  | 16.678 | <0.001 |
| L.IFG → R.ATL | 0.070 | 0.034 | 10.025 | <0.001 |  | 0.015 | 0.002 | 39.115 | <0.001 |  | 8.015 | <0.001 |
| L.IFG → R.IFG | 0.095 | 0.028 | 15.971 | <0.001 |  | 0.014 | 0.001 | 48.892 | <0.001 |  | 13.239 | <0.001 |
| L.IFG → L.pMTG | 0.093 | 0.036 | 12.483 | <0.001 |  | 0.015 | 0.002 | 44.159 | <0.001 |  | 10.359 | <0.001 |
| L.IFG → R.pMTG | 0.100 | 0.038 | 12.811 | <0.001 |  | 0.016 | 0.002 | 48.533 | <0.001 |  | 10.825 | <0.001 |
| R.IFG→ L.ATL | 0.139 | 0.047 | 14.326 | <0.001 |  | 0.014 | 0.002 | 34.124 | <0.001 |  | 12.784 | <0.001 |
| R.IFG → R.ATL | 0.083 | 0.029 | 13.799 | <0.001 |  | 0.014 | 0.002 | 41.184 | <0.001 |  | 11.123 | <0.001 |
| R.IFG → L.IFG | 0.094 | 0.030 | 15.105 | <0.001 |  | 0.014 | 0.001 | 48.779 | <0.001 |  | 12.888 | <0.001 |
| R.IFG → L.pMTG | 0.085 | 0.034 | 11.999 | <0.001 |  | 0.015 | 0.002 | 48.345 | <0.001 |  | 9.774 | <0.001 |
| R.IFG → R.pMTG | 0.077 | 0.059 | 6.296 | <0.001 |  | 0.016 | 0.001 | 50.755 | <0.001 |  | 4.975 | <0.001 |
| L.pMTG → L.ATL | 0.135 | 0.045 | 14.547 | <0.001 |  | 0.016 | 0.002 | 31.618 | <0.001 |  | 12.419 | <0.001 |
| L.pMTG → R.ATL | 0.090 | 0.054 | 7.901 | <0.001 |  | 0.016 | 0.002 | 39.670 | <0.001 |  | 6.481 | <0.001 |
| L.pMTG → L.IFG | 0.101 | 0.028 | 16.961 | <0.001 |  | 0.015 | 0.002 | 45.684 | <0.001 |  | 14.778 | <0.001 |
| L.pMTG → R.pMTG | 0.083 | 0.048 | 8.256 | <0.001 |  | 0.015 | 0.001 | 50.875 | <0.001 |  | 6.706 | <0.001 |
| L.pMTG → R.pMTG | 0.096 | 0.068 | 6.781 | <0.001 |  | 0.017 | 0.002 | 53.068 | <0.001 |  | 5.595 | <0.001 |
| R.pMTG→ L.ATL | 0.123 | 0.039 | 15.008 | <0.001 |  | 0.016 | 0.002 | 34.220 | <0.001 |  | 12.575 | <0.001 |
| R.pMTG → R.ATL | 0.086 | 0.051 | 8.116 | <0.001 |  | 0.016 | 0.002 | 41.485 | <0.001 |  | 6.533 | <0.001 |
| R.pMTG → L.IFG | 0.113 | 0.052 | 10.330 | <0.001 |  | 0.016 | 0.002 | 48.362 | <0.001 |  | 8.966 | <0.001 |
| R.pMTG → R.IFG | 0.077 | 0.070 | 5.258 | <0.001 |  | 0.016 | 0.001 | 51.017 | <0.001 |  | 4.185 | <0.001 |
| R.pMTG → R.pMTG | 0.103 | 0.075 | 6.610 | <0.001 |  | 0.017 | 0.002 | 50.808 | <0.001 |  | 5.547 | <0.001 |

Table S2. Results of intrinsic connectivity. L = left hemisphere, R = right hemisphere

**Supplementary Table 3**

| Semantic task | ATL stimulation | | | |  | Ventricle (control) stimulation | | | |  | ATL stimulation vs. Ventricle stimulation | |
| --- | --- | --- | --- | --- | --- | --- | --- | --- | --- | --- | --- | --- |
| Parameter |  |  |  |  |  |  |  |  |  |  |  | |
| Modulatory connectivity | Mean | SD | t | p |  | Mean | SD | t | p |  | t | p |
| L.ATL → R.ATL | -0.677 | 1.276 | -2.545 | 0.018 |  | 0.006 | 0.005 | 5.739 | <0.001 |  | -2.566 | 0.018 |
| L.ATL → L.IFG | -0.488 | 0.657 | -3.561 | 0.002 |  | 0.004 | 0.002 | 8.395 | <0.001 |  | -3.588 | 0.002 |
| L.ATL → R.IFG | -0.473 | 0.727 | -3.120 | 0.005 |  | 0.004 | 0.002 | 9.663 | <0.001 |  | -3.145 | 0.005 |
| L.ATL → L.pMTG | -1.707 | 0.845 | -9.686 | <0.001 |  | 0.006 | 0.003 | 10.913 | <0.001 |  | -9.721 | <0.001 |
| L.ATL → R.pMTG | -2.389 | 0.983 | -11.657 | <0.001 |  | 0.007 | 0.004 | 7.160 | <0.001 |  | -11.685 | <0.001 |
| R.ATL → L.ATL | 1.159 | 0.668 | 8.316 | <0.001 |  | 0.007 | 0.006 | 5.385 | <0.001 |  | 8.226 | <0.001 |
| R.ATL → L.IFG | -0.134 | 0.236 | -2.715 | 0.013 |  | 0.004 | 0.002 | 8.436 | <0.001 |  | -2.805 | 0.01 |
| R.ATL → R.IFG | -0.125 | 0.205 | -2.918 | 0.008 |  | 0.004 | 0.002 | 10.918 | <0.001 |  | -3.020 | 0.006 |
| R.ATL → L.pMTG | -0.309 | 0.259 | -5.704 | <0.001 |  | 0.006 | 0.002 | 12.545 | <0.001 |  | -5.820 | <0.001 |
| R.ATL → R.pMTG | -0.382 | 0.202 | -9.084 | <0.001 |  | 0.007 | 0.005 | 6.937 | <0.001 |  | -9.214 | <0.001 |
| L.IFG → L.ATL | 1.329 | 0.445 | 14.333 | <0.001 |  | 0.004 | 0.003 | 7.442 | <0.001 |  | 14.269 | <0.001 |
| L.IFG → R.ATL | -0.220 | 0.202 | -5.221 | <0.001 |  | 0.004 | 0.002 | 11.088 | <0.001 |  | -5.314 | <0.001 |
| L.IFG → R.IFG | -0.186 | 0.165 | -5.397 | <0.001 |  | 0.004 | 0.001 | 12.879 | <0.001 |  | -5.509 | <0.001 |
| L.IFG → L.pMTG | -0.323 | 0.152 | -10.180 | <0.001 |  | 0.005 | 0.002 | 14.317 | <0.001 |  | -10.320 | <0.001 |
| L.IFG → R.pMTG | -0.393 | 0.151 | -12.484 | <0.001 |  | 0.006 | 0.003 | 8.792 | <0.001 |  | -12.630 | <0.001 |
| R.IFG→ L.ATL | 1.264 | 0.426 | 14.211 | <0.001 |  | 0.004 | 0.002 | 8.104 | <0.001 |  | 14.157 | <0.001 |
| R.IFG → R.ATL | -0.226 | 0.180 | -6.031 | <0.001 |  | 0.004 | 0.001 | 11.483 | <0.001 |  | -6.235 | <0.001 |
| R.IFG → L.IFG | -0.207 | 0.094 | -10.544 | <0.001 |  | 0.004 | 0.001 | 14.800 | <0.001 |  | -10.708 | <0.001 |
| R.IFG → L.pMTG | -0.304 | 0.138 | -10.525 | <0.001 |  | 0.005 | 0.002 | 14.629 | <0.001 |  | -10.666 | <0.001 |
| R.IFG → R.pMTG | -0.337 | 0.172 | -9.376 | <0.001 |  | 0.006 | 0.003 | 9.526 | <0.001 |  | -9.498 | <0.001 |
| L.pMTG → L.ATL | 0.836 | 0.347 | 11.557 | <0.001 |  | 0.006 | 0.004 | 7.523 | <0.001 |  | 11.428 | <0.001 |
| L.pMTG → R.ATL | -0.074 | 0.270 | -1.312 | 0.203 |  | 0.006 | 0.002 | 12.679 | <0.001 |  | -1.414 | 0.171 |
| L.pMTG → L.IFG | 0.014 | 0.178 | 0.387 | 0.702 |  | 0.006 | 0.002 | 12.837 | <0.001 |  | 0.229 | 0.821 |
| L.pMTG → R.pMTG | -0.134 | 0.317 | -2.030 | 0.055 |  | 0.005 | 0.002 | 14.125 | <0.001 |  | -2.103 | 0.047 |
| L.pMTG → R.pMTG | -0.489 | 0.169 | -13.891 | <0.001 |  | 0.009 | 0.004 | 10.340 | <0.001 |  | -14.075 | <0.001 |
| R.pMTG→ L.ATL | 0.534 | 0.281 | 9.099 | <0.001 |  | 0.008 | 0.002 | 15.766 | <0.001 |  | 8.940 | <0.001 |
| R.pMTG → R.ATL | -0.082 | 0.325 | -1.206 | 0.24 |  | 0.008 | 0.003 | 14.160 | <0.001 |  | -1.313 | 0.203 |
| R.pMTG → L.IFG | 0.133 | 0.331 | 1.919 | 0.068 |  | 0.007 | 0.002 | 14.101 | <0.001 |  | 1.818 | 0.083 |
| R.pMTG → R.IFG | -0.179 | 0.412 | -2.089 | 0.048 |  | 0.007 | 0.002 | 15.356 | <0.001 |  | -2.163 | 0.042 |
| R.pMTG → R.pMTG | -0.283 | 0.385 | -3.519 | 0.002 |  | 0.009 | 0.003 | 17.765 | <0.001 |  | -3.632 | 0.001 |
| Driving input |  |  |  |  |  |  |  |  |  |  |  |  |
| tbTUS in the L.ATL | -3.589 | 0.240 | -71.784 | <0.001 |  |  |  |  |  |  |  |  |

Table S3. Results of modulatory connectivity during semantic processing. L = left hemisphere, R = right hemisphere

**Supplementary Table 4**

|  | Ventricle tbTUS | ATL tbTUS | χ2 | p |
| --- | --- | --- | --- | --- |
| Headache | 0 | 0 | − | − |
| Unusual feeling on the skin | 2 | 3 | 0.178 | 0.673 |
| Neck pain | 2 | 1 | 0.407 | 0.524 |
| Tingling | 3 | 3 | 0.003 | 0.953 |
| Itchiness | 0 | 0 | − | − |
| Difficulty paying attention | 3 | 2 | 0.278 | 0.598 |
| Unusual feelings, attitude, emotions | 0 | 0 | − | − |
| Sleepiness | 5 | 4 | 3.374 | 0.497 |
| Change in hearing | 1 | 0 | 1.069 | 0.301 |
| Nausea/stick to stomach | 0 | 0 | − | − |
| Dizziness | 2 | 0 | 2.188 | 0.139 |
| Anxious/worried/nervous | 0 | 0 | − | − |
| Forgetful | 0 | 0 | − | − |
| Difficulty with your balance | 1 | 1 | 0.001 | 0.974 |
| Other | 17 | 19 | 4.326 | 0.115 |
| (heard faint clicking/tapping sound during the stimulation) |  |  |  |  |

Table S4. The results of aversive questionnaires. The number was the number of participants.
